# Overexpression of osmosensitive Ca^2+^-activated channel TMEM63B promotes migration in HEK293T cells

**DOI:** 10.1101/626010

**Authors:** Marta C. Marques, Inês S. Albuquerque, Sandra H. Vaz, Gonçalo J. L. Bernardes

## Abstract

The recent discovery of the osmosensitive calcium (Ca^2+^) channel OSCA has revealed the potential mechanism by which plant cells sense diverse stimuli. Osmosensory transporters and mechanosensitive channels can detect and respond to osmotic shifts that play an important role in active cell homeostasis. TMEM63 family of proteins are described as the closest homologues of OSCAs. Here, we characterize TMEM63B, a mammalian homologue of OSCAs, recently classified as mechanosensitive. In HEK293T cells TMEM63B localizes to the plasma membrane and is associated to F-actin. This Ca^2+^-activated channel specifically induces Ca^2+^ influx across the membrane in response to extracellular Ca^2+^ concentration and hyperosmolarity. In addition, overexpression of TMEM63B in HEK293T cells significantly enhanced cell migration and wound healing. The link between Ca^2+^ osmosensitivity and cell migration might help to establish TMEM63B’s pathogenesis, for example in cancer in which it is frequently overexpressed.

## INTRODUCTION

More than 1000 families of transport proteins have been already recognized according to the IUBMB-approved Transporter Classification Database.[1] Recently, a new Anoctamin (ANO) superfamily of Ca^2+^-activated ion channels has been identified, which includes anoctamins (lipid scramblases), transmembrane channels (TMC) and Ca^2+^-permeable stress-gated cation channel (CSC) families.[2] Within this superfamily is the CSC-like family TMEM63, which shares the same topologies as the CSC family.[3] The lack of information about the molecular nature of the TMEM63 family encouraged us to investigate these ion channels.

Numerous proteins, such as *At*OSCA1.1 and *At*CSC1-OSCA1.2, found in *Arabidopsis thaliana* have been identified as mechanosensitive and were structurally characterized.[4-6] In mammals, some cation channels have been proposed to mediate osmosensory transduction through proportional modulation of their probability of opening during changes in fluid osmolality, but the molecular identity of these channels remains unknown.[7] Among these are the OSCAs orthologues, the TMEM63 family, proposed as likely candidates for the mammalian central osmosensory transduction pathway.[8] Interestingly, a recent study has linked OSCA/TMEM63 A and B channels to a mechanosensory role.[9] Investigation of the conserved domain architecture among these transporters revealed the existence of orthologs present in various taxonomic groups, such as fungi, green algae, plants, birds and mammals, which suggests the functional conservation of this family throughout higher eukaryotes. With the availability of sequences of various ANO superfamily members and the recently reported 3D structures of some OSCA proteins in *Arabidopsis thaliana*^4–6^ it is possible to get an insight into the structural aspects of TMEM63B domains involved in the ion translocation across the membrane.

## Results & Discussion

### TMEM63B shares 3D homology with OSCA1

By using BLAST[10] software we performed similarity searches with *Mus musculus* TMEM63B (*Mm*TMEM63B) as query protein (Uniprot: Q3TWI9) against protein structures in the Protein Data Bank (PDB).[11] The protein with the highest alignment score (**Table S1**) corresponds to *A. thaliana* CSC1 (OSCA1.2), followed by *At*OSCAs 3.1 and 1.1. The bioinformatic analysis of *Mm*TMEM63B amino acid sequence showed the presence of three different domains, a cytoplasmic PHM7 cyt domain and a N-and C-terminal Ca^2+^-dependent channel domain (RSN1), which consists of a total of 11 transmembrane (TM) predicted regions (**Figure S1**). We selected ten representative members of the ANO superfamily and performed a multiple sequence alignment with the MAFFT 7 program[12] (**Figure S2**) to identify functionally and structurally conserved residues and built a phylogenetic tree (**Figure 1A**). The structure and function of a protein relies on coordinated interactions among its residues.[13] Therefore, the relationship of other structural features of TMEM63B residues could be done through the identification of residues that co-vary with each other during evolution. We have found several residues with higher connectivity in the network (**Figure 1B** and **Table S2**), such as P492, P451, S508, D538, N177, K517, E345 and A332, which are also residues with a high degree of conservation. To produce a model of TMEM63B tertiary structure, *At*CSC1-OSCA1.2 3D structure was selected as a template for homology modelling by using the SWISS-MODEL server.[14] OSCA1.2 structure and predicted TMEM63B 3D model appear to share a similar 3D-fold (**Figure 1C**). In the obtained homology model, it is possible to observe several helices and a smaller number of strands that are localized within conserved regions of OSCA proteins, in agreement with the bioinformatic predictions. The computed degree of conservation for each residue was then mapped onto the predicted model of TMEM63B (**Figures 1C, 1D** and **Table S3**). The conserved patches essentially fall in TM1, TM4-TM7 and TM9-TM10. The cytosolic domain of OSCAs comprise a RNA recognition motif (RRM)[4, 6] that is also present in TMEM63B (**Figure S3A**). Based on an intensive search for similar sequences by using the SWISS-MODEL server[14] and PDB,[11] we identified three consensus sequences (see **Figure S3B**) that are conserved in TMEM63A and B and a family of RNA binding proteins, the heterogeneous Nuclear Ribonucleoprotein A1 (hnRNP A1, PDB:4yoe) and U6 snRNP-binding protein (PDB:2do4). However, the capability of TMEM63B to bind RNA remains undetermined and should be investigated in future studies.

**Figure 1.**
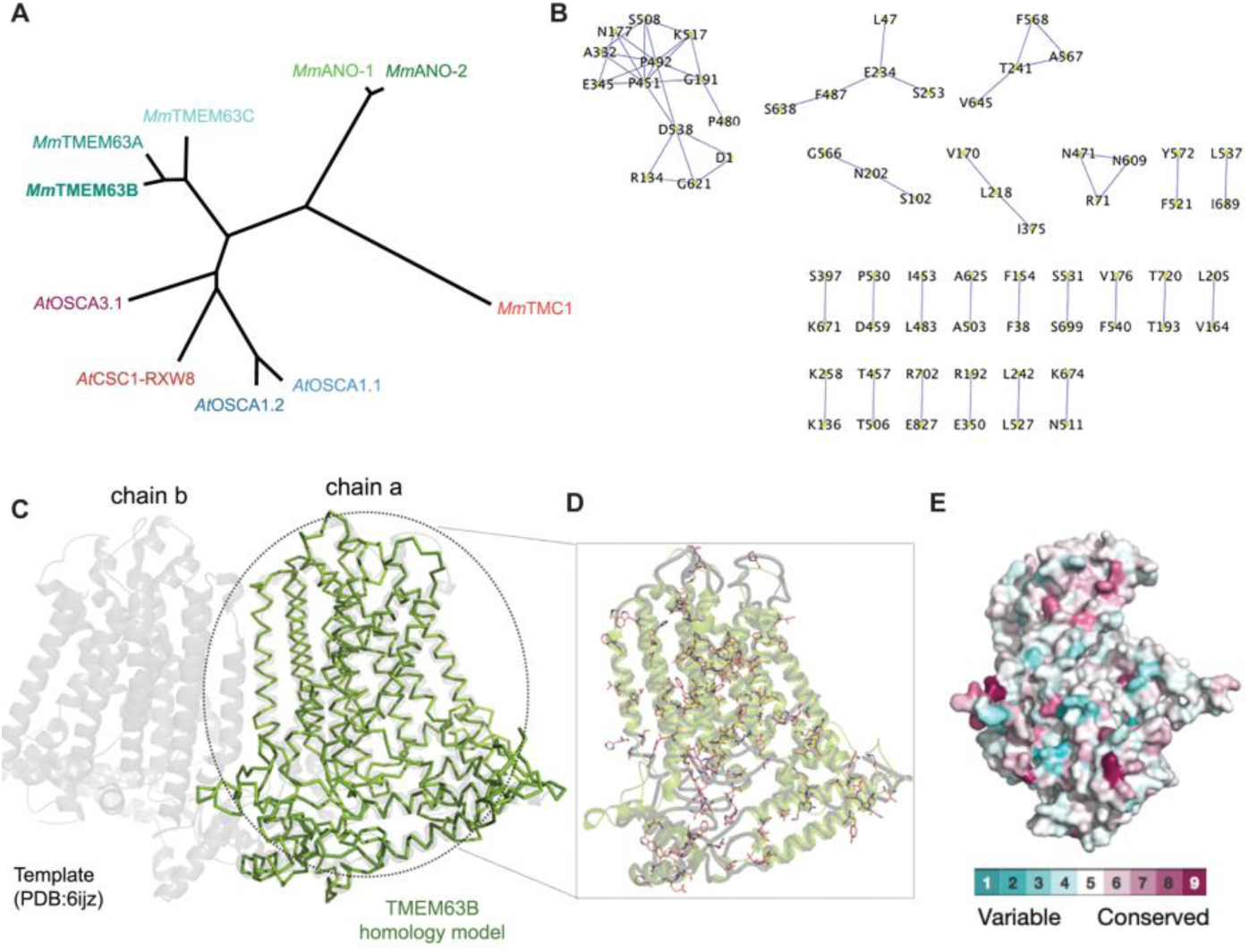
TMEM63B is an evolutionary conserved protein that shares 3D homology with OSCA1. **A.** Bayesian tree of proteins with similarity to TMEM63 family (teal clade), OSCA1 (blue clade), CSC1 (magenta clade), OSCA3 (purple clade), TMC (red clade) and TMEM16 (green clade), all from *M. musculus*. Tree was generated in DrawTree.[15] **B.** Co-evolution residue network analysis of ten homologous proteins of *M. musculus* TMEM63B. The analysis was performed with CAPS[16] software to study coevolving amino acids. Co-evolving amino acid pairs were defined in terms of their statistical support (defined as bootstrap values); only pairs with bootstrap values above 0.8 were used for construction of networks with Cytoscape software.[17] **C.** 3D homology model of the mouse TMEM63B protein in green, represented as a ribbon built with OSCA 1.2 as template (PDB:6ijz), represented in grey as cartoon by using Pymol.[18] **D.** Partial representation of *Mm*TMEM63B (green) with the predicted 3D location of the conserved residues (pink element sticks). **E.** ConSurf analysis for the homology model built with Modeller[19] for TMEM63B. The 3D structure is rendered as surface and color coded by its conservation grade by using the color-coding bar shown in the figure, with turquoise-through bordeaux that indicates the variable to conserved residues. The analysis was carried out with MAFFT multiple sequence alignment[12] available and the figures were generated with the help of PyMOL[18] script output by ConSurf.[20]

### TMEM63B promotes [Ca^2+^]_i_ influx in response to extracellular [Ca^2+^] and hyperosmolarity

The putative ion conduction pore and the mechanosensitive features of OSCAs structures showed to be partially conserved in TMEM63B (**Figure S3B**). Based on this data and with the assumption that TMEM63 proteins could be osmosensitive calcium-activated channels, we decided to clone the TMEM63B gene from *M. musculus*, fused to enhanced green fluorescent protein (EGFP) and overexpress it in Human Embryonic Kidney 293 (HEK293T) cells. We confirmed the TMEM63B-EGFP construct expression (**Figure 2A**) and loaded cells with the calcium-responsive dye Fura-2AM. Stimulation with 2 mM of Ca^2+^ elicited an intracellular calcium increase (**Figure 2B**), significantly higher than the control (p < 0.0001), which suggests that TMEM63B is a calcium-activated channel. To provide further support for our hypothesis, we also challenged HEK293T cells transfected with TMEM63B-EGFP fusion protein with 150 mM of mannitol, to simulate hypertonic shock, and evaluated [Ca^2+^]_i_ oscillation (**Figure 2C**). The [Ca^2+^]_i_ increase induced by mannitol was significantly higher in cells transiently overexpressing TMEM63B-EGFP, than those expressing the empty vector (p < 0.0002) (**Figure 2C and D**). Exposing cells to a hypertonic solution is a form of mechanical stress that caused cells to retract (**Figure 2E, Supporting V1** and **V2**). Still, the importance of choosing cells that lack endogenous mechanically activated (MA) channels to assess the mechanotransduction properties is crucial. It is known that HEK293T cells possess the MA channels Piezo and NOMPC, that can also induce MA currents.[21, 22] Yet, it is clear that the transfected cells react to the mannitol shock, still this response is moderate relative to stimulation with [Ca^2+^] (**Figure 2F, 2G** and **Supporting V2**). We also tested a higher mannitol concentration (300 mM) but the results were not significantly different (data not shown). Strikingly, a recent study confirmed *Mm*TMEM63B as mechanosensitive, inducing stretch-activated currents when expressed in naïve cells.[9] Systemic osmoregulation is a crucial process whereby changes in plasma osmolality, detected by osmoreceptors, modulate renal function and stabilize the tonicity and volume of the extracellular fluid.[7, 23] Because TMEM63B shows distinct membrane expression in several cell types, being widely expressed in the kidneys (renal tubules), epididymis, lungs and tonsil,[24] it is valid to assume that this channel might contribute to osmosensitive Ca^2+^ entry into these tissues.

**Figure 2.**
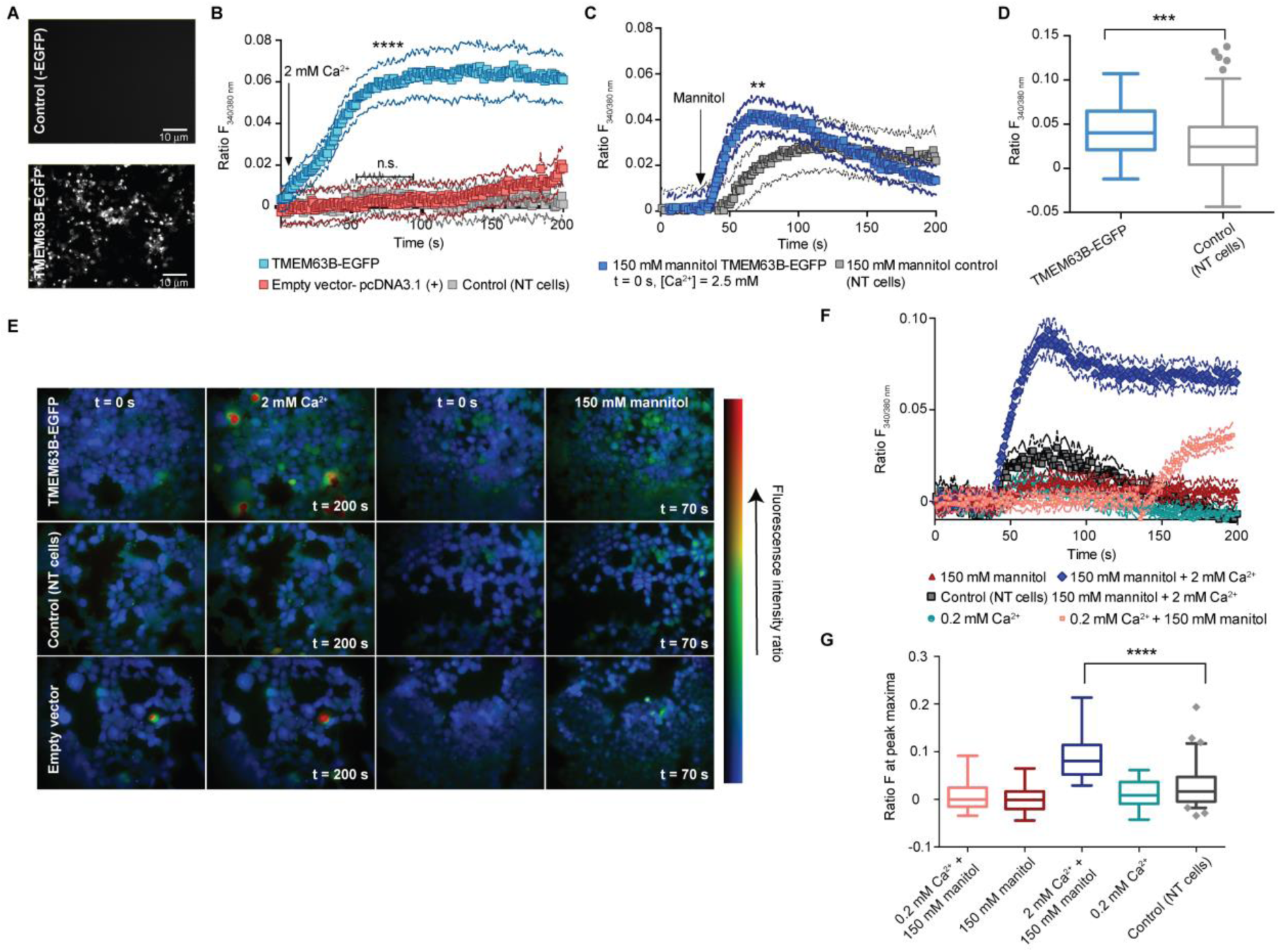
TMEM63B promotes [Ca^2+^]_i_ influx across the plasma membrane in response to extracellular [Ca^2+^] and hyperosmolarity. **A.** Representative fluorescent images showing the HEK293T cells either non transfected or transfected and showing EGFP positive signal. Scale-bars represent 10 µm. **B.** Increase in calcium influx induced by extracellular Ca^2+^ (2 mM) in EGFP-positive HEK293T cells. For quantification, the strongest responding cells were analyzed in each well (20–30 cells, n=3) and normalized to the baseline. **** Mann-Whitney of unpaired t test data: significant difference between empty vector and TMEM63B-EGFP (p < 0.0001) after peak determination by the analysis of the area under the curve. No significant difference between control and NT. **C**. Increase in calcium influx induced by extracellular mannitol (150 mM) in EGFP-positive HEK293T cells. For quantification, the strongest responding cells were analyzed in each well (20-30 cells, n=3) and normalized to the baseline. ** Mann-Whitney of unpaired t test data between TMEM63B-EGFP and control (NT cells) (p < 0.0002) at peak maxima. **D.** A box plot of peaks at 70 s post mannitol stimulation of HEK293T cells (control: NT cells). ***Mann-Whitney of unpaired t test data: significant difference between NT cells and TMEM63B-EGFP (p < 0.0001). **E.** Calcium imaging of HEK293T cells transfected with the mouse TMEM63B-EGFP fusion protein or a pcDNA3.1(+) empty vector and non transfected, corrected for baseline level before and after calcium (2 mM) or mannitol stimuli (150 mM) (color code – blue = low calcium level; green = intermediate; yellow = medium high; red = high). **F.** Calcium influx induced by extracellular Ca^2+^ (0.2 and 2 mM) with a constant mannitol concentration of 150 mM. For quantification, the strongest responding cells were analyzed in each well (n=30 cells) and normalized to the baseline. For the condition where we used 0.2 mM Ca^2+^ and 150 mM mannitol (salmon symbols) we applied a second mannitol shock at 120 s. **G.** A box plot of peaks after either calcium or mannitol stimulation at peak maxima. ****Mann-Whitney of unpaired t test data: significant difference between NT cells and TMEM63B-EGFP at 150 mM mannitol and 2 mM of Ca^2+^ stimulation (p <0.0001). NT cells, non-transfected cells. Values represent mean ± s.e.m..

### TMEM63B enhances cell migration in a calcium-dependent manner

We also determined the localization of TMEM63B-EGFP (**Figure 3A**) by confocal microscopy and the protein was found to be predominantly associated to the plasma membrane, cortical F-actin, but not vinculin (**Figure 3B** and **S4**).

**Figure 3.**
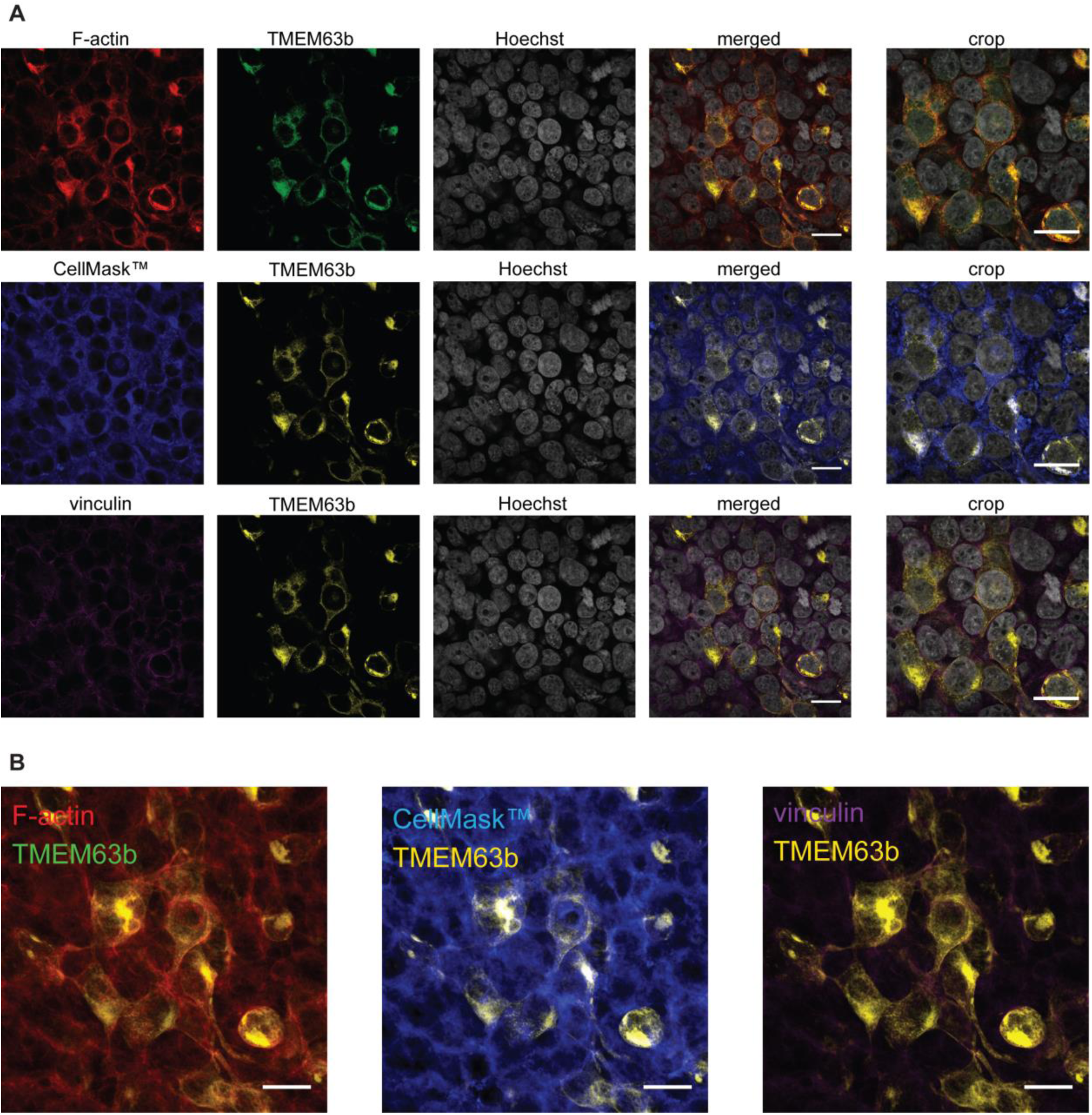
TMEM63B localizes to the membrane and co-localizes with cortical F-actin but, not with vinculin. **A.** Confocal microscopy images of 6 µm optical slices of fixed HEK293T cells transfected with TMEM63B-EGFP and stained with phalloidin (red), CellMask™ Plasma membrane Stain (blue), anti-vinculin (magenta), and Hoechst 33342 (grey). On the right panel, we show a crop of the merged channels, so as to further accent colocalization or lack thereof between these markers and *Mm*TMEM63B. *Mm*TMEM63B is shown in either green or yellow. **B.** Z-projection of the merged channels, corresponding to the same imaged fields shown above. Scale-bars represent 10 µm. Colocalization analysis (Pearson’s coefficient): TMEM63B *vs* vinculin r = 0.039; TMEM63B *vs* F-actin r = 0.701; TMEM63B *vs* cell membrane r = 0.393, showing partial colocalization with cell membrane and F-actin, but not with vinculin.

It is known that the actin network is involved in several cellular processes related to the control of dynamic cellular morphology, organelle organization, and motility in reaction to various chemical and mechanical signals.[25] Dysfunction in proteins in the actin and focal adhesion proteomes are associated with numerous severe diseases, such as muscular disorders and cancers. Moreover, the intracellular Ca^2+^ has a great impact on the migration machinery of healthy, tumor and stromal cells.[26-28] Based on this combined information, the role of TMEM63B in HEK293T cells was speculated to regulate cell migration. To demonstrate this, we transiently overexpressed TMEM63B-EGFP fusion protein (**Figure 4A**) in HEK293T cells, that express a low level of TMEM63B (**Figure 4B**). We first applied a scratch and then evaluated by microscopy the cell wound closure and the ability of the HEK293T cells to migrate and subsequently close the wound made in a confluent plate of cells, after 24 h(see **Figure 4C**). Based on the width of the wound, we calculated the percentage of wound healing (**Figure 4D**). The tendency of HEK293T cells to detach from the plate immediately after the wound is made was prevented by coating with 0.1% gelatin. Remarkably, our data indicated that overexpression of TMEM63B-EGFP significantly increased migration of HEK293T cells relative to control cells (p < 0.02). Cell migration is a central component of the metastatic cascade, which requires a concerted action of ion channels and transporters, cytoskeletal elements and signaling. The migration cycle demand spatially synchronized changes of the actin cytoskeleton[28, 29] in which TMEM63B might play an important role. We also evaluated the effect in wound healing of the cell permeant Ca^2+^ chelator bapta-AM. Treatment with the Ca^2+^ chelating agent caused some degree of cell detachment. Incubation of the cells with bapta-AM reduced the percentage of wound healing both for TMEM63B-EGFP overexpressing cells and control (**Figure 4E** and **4F**), which suggests the existence of a Ca^2+^-dependent migration process. Studies have reported that several ion channels contribute to a variety of basic cell processes such as proliferation, adhesion, migration and invasion by inducing local volume changes and/or by modulating Ca^2+^ influx, crucial for carcinogenesis and cancer development.[30, 31] Interestingly, TMEM63B was one of the mRNAs, among a few other, which expression appears to be downregulated by miR-199a-5p in mouse keratinocytes,[32] a small non-coding RNA molecule. These molecules mediate diverse biological cellular processes through regulatory pathways, targeting genes through translational repression or mRNA degradation.[33] Previous studies have also implicated the same miR-199a-5p gene targets in cell proliferation and migration in cancer cells. Intrigued by these data, we also evaluated TMEM63B mRNA expression in different cancer cell lines (lung, hepatocyte and ductal carcinomas and breast adenocarcinoma). We found a differential gene expression among the different cell lines, being the mRNA expression level of T47-D cell line (ductal carcinoma) more than two-fold higher than HEK293T cells, as shown in **Figure 4G**.

**Figure 4.**
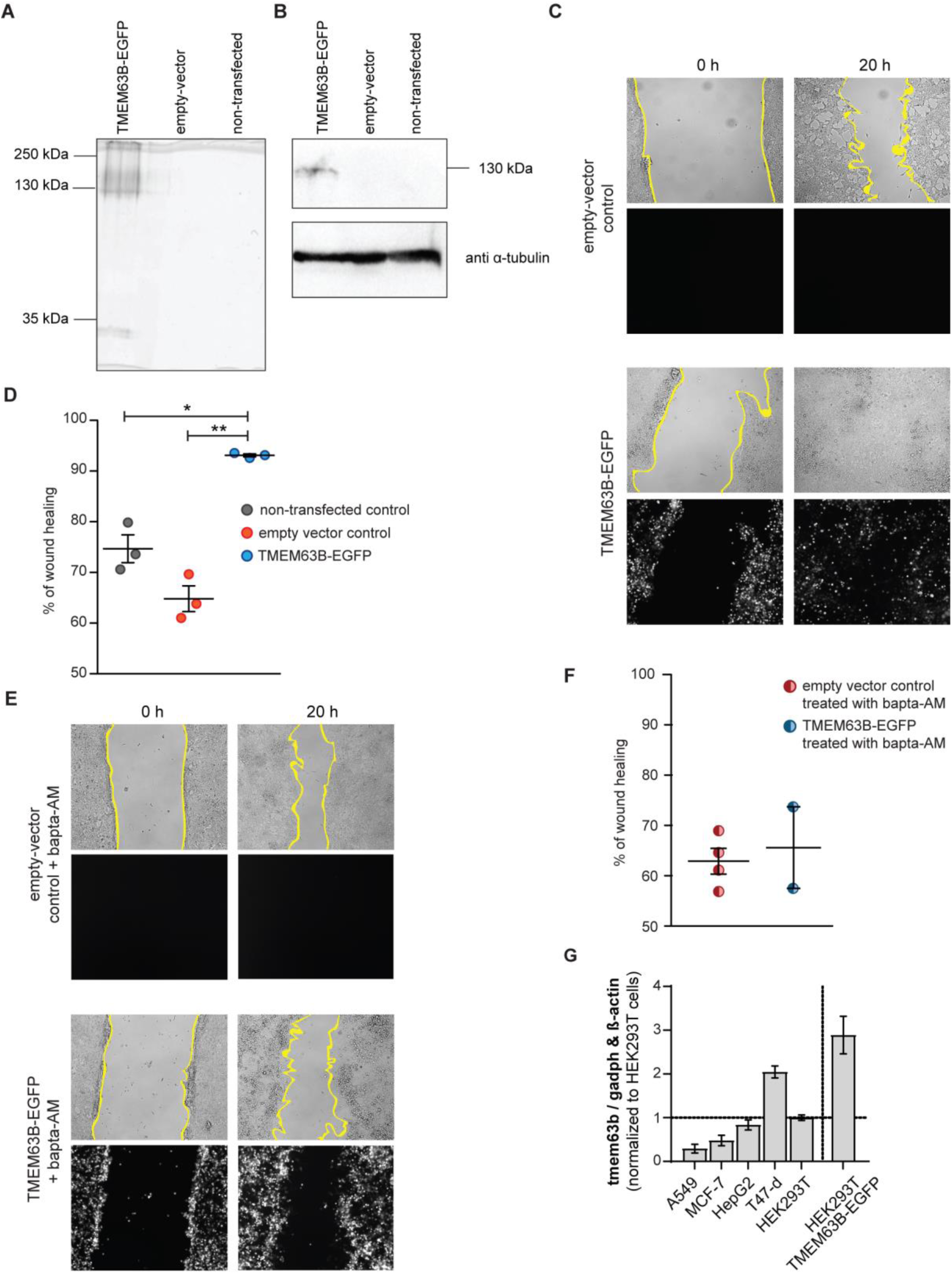
TMEM63B overexpression in HEK293T enhances cell migration in a calcium-dependent manner. **A.** Whole HEK293T cell lysates were assayed for 10 % SDS-gel with fluorescence imaging with a Typhoon fluorescence scanner (excitation: 488 nm laser) and **B.** western blot. Cell lysates from controls (empty vector and non-transfected cells) and *Mm*TMEM63B-EGFP-cells, were quantified and 50 µg from each protein extract were loaded on a 10% SDS-PAGE gel, as indicated. The SDS gel was imaged on a Typhoon scanner, and then transferred to a PVDF membrane. **C.** Representative transmitted-light widefield microscopy images of cells transfected with an empty vector or with *Mm*TMEM63B-EGFP. Images were taken as soon as the scratch was performed (0 h) or, at the same position, after 20 h. **D.** Quantification of the percentage of wound healing in a 20 h scratch-assay. Bars indicate average and error bars indicate standard error of the mean. **E.** Representative transmitted-light widefield microscopy images of cells transfected with an empty vector or with *Mm*TMEM63B-EGFP, and incubated with bapta-AM, a known Ca^2+^-chelator. Images were taken as soon as the scratch was performed (0 h) or, at the same position, after 20 h. **F.** Quantification of the percentage of wound healing in a 20 h scratch-assay when cells were treated with bapta-AM. Statistical analysis was performed with a two-tailed unpaired t-test with Welch’s correction (** p<0.02 and * p<0.05). **G.** Quantification of mRNA levels of hTMEM63B in human tumor cell lines by quantitative real-time PCR. Data shown as ΔΔCT of tmem63b to house-keeping genes gadph and ß-actin, normalized to ΔCT for HEK293T cells.

## Conclusions

In summary, our results indicate that TMEM63B protein belongs to an osmosensitive ion channel family, which is conserved across eukaryotes, and has similar architecture to the OSCA family. We show that expression of TMEM63B in HEK293T cells promotes [Ca^2+^]_i_ influx across the plasma membrane in response to extracellular [Ca^2+^] and hyperosmolarity. Moreover, overexpression of TMEM63B-EGFP enhances cell migration in HEK293T cells. These results directly demonstrate that TMEM63B is linked to Ca^2+^ signaling on which cell migration is dependent, which suggests for the first time such a functional role for TMEM63B. We are now focusing on the identification of the molecular mechanisms by which TMEM63B overexpression affects cell migration. Furthermore, we found an increase in TMEM63B mRNA expression in the ductal carcinoma T47-D cell line, and we are conducting studies to clarify the potential role(s) of TMEM63B as a cancer biomarker.

## Method details

### Sequence alignment tools

Sequence data were recovered from the National Center for Biotechnology Information website (https://www.ncbi.nlm.nih.gov). IDs with 6 characters are UniProt Ids. The sequences of MmTMEM63B (UniProt Accession: Q3TWI9), MmTMEM63A (UniProt Accession: Q91YT8), MmTMEM63C (UniProt Accession: Q8CBX0), MmANO-1 (TMEM16A, UniProt Accession: Q8BHY3), and MmANO-2 (TMEM16B, UniProt Accession: Q8CFW1) and MmTMC1 (UniProt Accession: Q8R4P5), AtOSCA1.1 (UniProt Accession: Q9XEA1), AtCSC1-OSCA1.2 (UniProt Accession: Q5XEZ5), AtOSCA3.1 (UniProt Accession: A0A097NUQ9) and AtRXW8 (UniProt Accession: F4IBD7). Multiple alignments were performed using MAFFT 7,^6^ edited with Jalview.^7^ To further investigate phylogenetic relationships, we have compared the aligned sequences from all ten representatives of ANO Superfamily. A phylogenetic tree based on the multiple alignment generated was constructed through Bayesian phylogenetic inference using Phylo.io^13^. Tree was generated in DrawTree 3.67^14^ and illustrates the relationship between members of the CSC family across CSC1 (OSCA1) from *A. thaliana* (blue clade), CSC1-like (purple clade), TMEM63 (teal clade), TMC (red clade) and TMEM16 (green clade), all from *M. Musculus*.

### Bioinformatics analysis of mouse TMEM63B amino acid sequence

The sequence of *Mm*TMEM63B was used as query for the identification of conserved domains and secondary structure. The conserved domains of the sequences were identified from NCBI Conserved Domains search.^15^ The secondary structure of the domains in TMEM63B protein were predicted using JPred prediction software.^2^ In order to elaborate on physiological role and structural details of this functional domains of TMEM63B in mammals, we have analyzed the proteins that contain one or more of the domains present in the query sequence, using the Conserved Domain Architecture Retrieval Tool (CDART)^1^. The comparative gene analysis with *Arabidopsis thaliana* (*At*) identified the orthologous groups DUF221 and CSC1-like ERD4 genes and their association with calcium transport and stress by dehydration. To further investigate this, we have compared ten sequences representative from the Anoctamin Superfamily of Ca^2+^-activated ion channels and lipid scramblases. *M. musculus* TMEM63B protein analysis showing the results of binding site, solvent accessibility and protein disorder predictions was performed using PredictProtein.^3^ The secondary structure predictions of TMEM63B were obtained using the CFSSP: Chou and Fasman Secondary Structure Prediction server.^16^

We performed an additional intra-protein coevolution analysis, using the multiple alignment generated for the 10 homologues as input, using a co-evolution method implemented in the programme CAPS.^10^ For this analysis we used a threshold α value of 0.01 and a bootstrap value of 0.8. Amino acids pairs with bootstrap values above 0.8 were used to construct networks using the Cytoscape software.^17^

### Protein structure homology and modelling

It was not feasible to find a protein with available structures in PDB^9^, with similarity to the entire TMEM63B. We used the amino acid sequence of the *M. musculus* TMEM63B and subjected to BLASTp^8^ search against proteins with known 3D structure^9^ to find out the protein sharing highest alignment scores. Three different proteins with similarity to the *Mm*TMEM63B produced significant alignment results, which have available structures in public databases. Further, *Mm*TMEM63B sequence was used as query in the SWISSMODEL server^12^ to find proteins with known 3D structures sharing sequence and structure similarities to TMEM63B. After, we used *Mm*TMEM63B sequence to create a model in the SWISS‐MODEL server^12^.The protein used as a template for the homology modelling of TMEM63B was the *At*CSC1-OSCA1.2 (PDB: 6ijz). To assess the structural quality of the obtained homology model, the generated structure was further assessed using Molprobity.^11^ The degree of conservation was obtained from the amino acid multiple alignment which included ten sequences of representative members of the ANO Superfamily, by estimating site-specific evolutionary rates, and using the Bayesian tree generated in the first section as input, through the ConSurf server.^18^ Conservation score, grouped into a nine-color grade scale (where the most conserved and variable positions are marked in bordeaux and turquoise, respectively), was mapped onto the 3D structure of the generated model.

### Plasmid construction

Total RNA was extracted from a C57BL/6J mouse lung after mechanical homogenization in 2 ml of denaturing solution (4 M guanidine thiocyanate; 25 mM sodium citrate pH 7; 0,5% N-Lauroyllsarcosine; and 0,7% of freshly added β-mercaptoethanol, in Diethylpyrocarbonate (DEPC)-treated water). RNA was extracted using a Qiagen RNeasy minikit, according to the manufacturer“s instructions. The synthesis of the first-strand cDNA from the RNA templates was carried out using the Transcriptor First Strand cDNA Synthesis Kit (Roche). The amplification program was as follows: 25°C (10 min), 55°C (30 min) and 85°C (5 min). Mouse TMEM63B was then PCR amplified from the cDNA with the previously described^19^ primers:

- Forward GGTTGAATTCCACCATGCTGCCGTTCTTGCTGG,
- Reverse CCTTGGTACCTTACTGGTGAATCTCATTCTCTATGAGG.

For expression in mammalian cells TMEM63B was subcloned into a modified pcDNA3.1(+) vector, resulting in a plasmid coding for a C-terminal EGFP-FLAG −10x Histidine-tagged *Mm*TMEM63B fusion protein. The gene product was cloned with XbaI and XhoI and ligated with the T4 ligation enzyme (ThermoFisher) and sequence was checked by sequencing.

### Cell culture, transfection and treatment

All cell lines cells (ATCC) were cultured in DMEM containing 10% FBS (GIBCO, Grand Island, NY) and 1% pen-strep (Sigma-Aldrich, St. Louis, MO) in 10% CO2. HK293T cells were transfected with pcDNA3.1(+)-TMEM63B-EGFP and pcDNA3.1(+) plasmids. For calcium imaging, EGFP was used as a selective marker. Cells were seeded for 24 h before transfection and recorded between 48h and 72 h after transfection.

### Imaging of [Ca^2+^]_i_ inHEK293T cells

HEK293T cells were cultured to 30% confluency in DMEM containing 10% FBS (GIBCO, Grand Island, NY) and 1% pen-strep (Sigma-Aldrich, St. Louis, MO) in 10% CO2 at 37 °C. For transfection, Cells were seeded in µ-ibidi 8 well plates and overnight and transfected with TMEM63B-EGFP and pcDNA3.1(+) empty vector using Fugene HD Reagent (Promega) as described previously.^20^ The Ca^2+^ imaging assay was performed in the HEK293T cells 24h to 72 h after transfection using an inverted microscope with epifluorescent optics (Axiovert 135TV, Zeiss) and equipped with a high speed multiple excitation fluorimetric system (Lambda DG4, with a 175W Xenon arc lamp). Data were recorded by a CDD camera. Cells were loaded with the Ca^2+^ sensitive dye Fura-2 AM (5 µM; Life Technologies) for 45 minutes to perform the intracellular calcium measurements, as described^20^. Before the measurements the cell medium was replaced by warm standard buffer containing 119 mM NaCl, 2.5 mM Ca^2+^, 2 mM MgCl_2_, 5 mM KCl, 10 mM glucose, 25 mM HEPES, pH 7.4 (adjusted with NaOH), unless stated otherwise. For the hyperosmotic treatments, at 30 seconds the shock was applied by addition of 20 µL or 40 µL of a stock solution of 1.5 M of mannitol containing 129 mM NaCl, 2.5 mM Ca^2+^, 2 mM MgCl2, 5 mM KCl, 10 mM glucose, 25 mM HEPES, pH 7.4 (150 mM or 300 mM final mannitol final concentrations, respectively). For Ca^2+^ activation, Fura 2-loaded HEK293T cells were incubated in a standard buffer containing 129 mM NaCl, 0.1mM Ca^2+^, 2 mM MgCl2, 5 mM KCl, 10 mM glucose, 25 mM HEPES, pH 7.4 and 1 µL of 500 mM Ca^2+^ was added to the cells (t=30s).

Fura-2 AM emission ratiometric images emissions from 340 nm and 380 nm excitation were recorded for 30 seconds in the beginning of the experiments. Images were collected and analyzed using MetaFluor Fluorescence Ratio Imaging Software (Molecular Devices). The statistical analysis was performed with GraphPad Prism. Experiments were carried out at room temperature. For further analysis, the 25 to 30 most responsive cells per image were selected manually, based on the increases in [Ca^2+^]_i_ (from highest to lowest). Experiments were performed at least in triplicate, except stated otherwise.

### SDS-PAGE and Western blot analysis

HEK293TT cells were transfected with TMEM63B-EGFP-FLAG-10xHis, with a negative control (empty vector) and non-transfected were cultured for 48 h. Cells were washed with PBS and lysed in lysis buffer (20 mM Tris-HCl pH 7.6, 300 mM NaCl, 1% Triton-X100, 1 mM EDTA, 5% glycerol, and protease inhibitor cocktail (Roche) and frozen at −20°C. The crude cell extracts from HEK293T cells were run on for 1.5h in a 10% SDS–PAGE gel and monitored by Amersham Typhoon Biomolecular Imager (emission filter 488 nm). Proteins were then transferred to PVDF membrane for 1h at constant 350 mA. Membranes were blocked with 5% milk powder in Tris-buffered saline with 0.1% Tween20 for 1 h and immunodetection was performed with an antibody for TMEM63B (Merck). Protein content was quantified by Bradford.

### TMEM63B–EGFP subcellular localization analysis

HEK293T cells transfected with an empty vector, transfected with TMEM63B-EGFP-FLAG-10xHis, or non-transfected were cultured for 48 h as described above, in 24 well-plates in which the bottom of the well was previously covered with glass round coverslips. Culture medium was then removed, cells were washed with PBS 1x and fixed with a 4% paraformaldehyde (PFA) solution for 10 minutes at room-temperature (RT). PFA was then removed, and cells were washed three times with PBS 1x before proceeding with the staining protocol. Cells were permeabilized at RT for 10 min with 0.1 % Triton-X in PBS 1x and then blocking was performed by incubating the cells 1 % BSA in permeabilizing solution (blocking solution), for 30 min, also at RT. Cells were stained with phalloidin conjugated with AlexaFluor^®^ 555 (Cat#: A34055, Thermofisher, USA) and mouse anti-vinculin by incubating in blocking solution for 1 h, in the dark at RT. Cells were then washed three times with PBS 1x, before incubating with an anti-mouse IgG H&L conjugated with AlexaFluor^®^ 594, for 1 h, in the dark, at RT. Cells were then washed with PBS 1x, and counterstained with Hoechst 33342 solution at 5 µg/ml for 10 min (Thermofisher, USA). Cells were also counterstained with CellMask™ Deep Red Plasma Membrane Stain, before a final washing step with PBS 1x. Coverslips were then removed and mounted with Fluormount-G™□ mounting medium in microscope appropriate glass slides. Coverslips were allowed to dry and kept in the dark until visualization in a Zeiss LSM 710 confocal microscope, equipped with a Plan-Apochromat 63x/1.40 Oil DIC objective, a Diode 405-30 laser (Hoechst 33342), an Argon 488 laser (GFP), an Argon 514 laser (AlexaFluor^®^ 555), a DPSS 561-10 laser (AlexaFluor^®^ 594) and a HeNe633 laser (Deep Red), configured according to the emission spectra as described in the Fluorescence Spectra Viewer (ThermoFisher Scientific, USA, online-tool). Images were then processed using ImageJ 2.0.0 (Fiji) and colocalization was quantified with the Jacop plugin.

### Cell migration (wound healing assay)

Twenty-four well plates were coated with gelatin 0.1% in phosphate buffered saline (PBS) for 30 minutes at 37 °C. 80,000 cells/well were seeded and grown for 48h. A scratch was performed with a P10 pipette tip, the medium was exchanged to DMEM supplemented with 2% FBS. After washing three times with PBS 1x, cells were imaged on a Zeiss Cell Observer, using the multistage position function, while incubating at 37°C and in controlled 5 % CO^2^ conditions. In some experiments HEK293T cells were incubated with bapta-AM (35 μM) or the control (0.14% DMSO) for 20 minutes before the scratch. Cells were returned to the incubator and the wounds were once again imaged at the same coordinates 24 h later. The wound areas were then quantified using ImageJ v2.0.0 (Fiji) software, and the MRI Wound Healing Tool. The percentage of wound healing in each well was calculated using the following formula: 100 – [(wound area at 24 h / wound area at 0 h) x 100].

### Quantification of hTMEM63b mRNA levels in human cell lines by quantitative real-time PCR

The quantitative real time PCR method was used to determine the levels of mRNA of human TMEM63B in different human cell lines. The samples were normalized to the mean of the internal controls GADPH and β-actin. Briefly, after trypsinization, cells from three confluent wells of a 24 well-plate were pooled, and total RNA was extracted using the NZY Total RNA Isolation Kit (NZYTech). From these samples, 1 µg of RNA was reverse transcribed using the NZY First-Strand cDNA Synthesis Kit (NZYTech) using random hexamers, according to the following manufacturer’s instructions. hTMEM63B cDNA was then quantified by quantitative real-time PCR, using the iTaq™ Universal SYBR^®^ Green Supermix (Bio-Rad) following the manufacturer’s instructions. The run was performed on 96 well-plate using a 7500Fast PCR System (Applied Biosystems™□). PCR was performed as followed: 2-min initial annealing at 50°C followed by 10-min denaturation at 95 °C, and then 50 cycles of amplification following 95 °C for 15 s denaturation, 60 °C for 1-min Annealing/Extension + Plate Read at 60°C. All reactions were performed on duplicate to ensure the reproducibility of the qPCR assay. Primers for amplification of specific hTMEM63B cDNA were synthesized by Sigma-Aldrich/Merck and are:

- Forward primer: 5’-TATCACCGCCATCATCCTGAAG-3’,
- Reverse primer: 5’-TGGAGAGGTGCTCCCAGTAG-3’.

## Data availability

Additional data and supporting figures and videos are available in the Extended View Content file.

## Acknowledgements

Funded under the Royal Society (URF\R\180019 to G.J.L.B.), FCT Portugal (iFCT IF/00624/2015 to G.J.L.B., Postdoctoral Fellowships SFRH/BPD/118731/2016 to M.C.M. and SFRH/BPD/81627/2011 to S.H.V., and Doctoral Studentship SFRH/BD/111556/2015 to M.I.A.) and ERC StG (GA No. 676832). The authors thank Dr Vikki Cantrill for her help with the editing of this manuscript.

## Author Contributions

M.C.M. conducted biochemical and cell biology studies. I.S. performed imaging while S.H.V. did Ca^2+^ flux assay measurements. M.C.M. and G.J.L.B. wrote the manuscript with contribution from the remaining authors. M.C.M. conceived the research. G.J.L.B. supervised research. All authors agreed on the final version of the manuscript.

